# Metabolic interplay between *Proteus mirabilis* and *Enterococcus faecalis* facilitates polymicrobial biofilm formation and invasive disease

**DOI:** 10.1101/2023.03.17.533237

**Authors:** Benjamin C. Hunt, Vitus Brix, Joseph Vath, Beryl L. Guterman, Steven M. Taddei, Brian S. Learman, Aimee L. Brauer, Shichen Shen, Jun Qu, Chelsie E. Armbruster

## Abstract

Polymicrobial biofilms play an important role in the development and pathogenesis of CAUTI. *Proteus mirabilis* and *Enterococcus faecalis* are common CAUTI pathogens that persistently co-colonize the catheterized urinary tract and form biofilms with increased biomass and antibiotic resistance. In this study, we uncover the metabolic interplay that drives biofilm enhancement and examine the contribution to CAUTI severity. Through compositional and proteomic biofilm analyses, we determined that the increase in biofilm biomass stems from an increase in the protein fraction of the polymicrobial biofilm matrix. We further observed an enrichment in proteins associated with ornithine and arginine metabolism in polymicrobial biofilms compared to single-species biofilms. We show that L-ornithine secretion by *E. faecalis* promotes arginine biosynthesis in *P. mirabilis,* and that disruption of this metabolic interplay abrogates the biofilm enhancement we see *in vitro* and leads to significant decreases in infection severity and dissemination in a murine CAUTI model.

## Introduction

Urinary tract infections (UTIs) are among the most common infections worldwide and account for approximately 40% of all nosocomial infections in the United States ^1–5^. UTIs are classified into two broad categories, uncomplicated and complicated UTI, dependent upon the presence of risk factors, disease severity, and location of infection ^1,3,5^. Urinary catheterization is a common procedure in healthcare settings with approximately 15-25% of patients at a general hospital acquiring a catheter at some point in their stay; the incidence of catheterization is even more frequent for the elderly, long-term care patients, and critically ill patients ^6–14^. Catheter insertion facilitates the development of bacterial colonization through a variety of means, including mechanical disruption, induction of inflammation, and by providing an ideal surface for bacterial attachment ^15–18^. Each day a urinary catheter is in place, there is a compounding 3-8% incidence of bacteriuria, and the majority of patients with long term catheterization (>28 days) will experience at least one symptomatic catheter-associated UTI (CAUTI) ^6,12,19–21^. CAUTI is one type of complicated UTI and is associated with high rate of treatment failure, increased patient morbidity and mortality, overuse of antibiotics, increased length of stay and hospital cost ^7,13,22,23^.

The epidemiology of CAUTI also differs from that of uncomplicated UTI; uncomplicated UTIs are most often caused by *Escherichia coli,* while complicated catheter-associated bacteriuria and CAUTI are caused by a more diverse range of pathogens in addition to *E. coli,* including *Proteus mirabilis, Enterococcus faecalis, Klebsiella spp., Pseudomonas aeruginosa,* and *Staphylococcus* species ^5,24–26^. Catheter-associated bacteriuria and CAUTI are frequently polymicrobial, which further complicate treatment efficacy and infection severity ^24,27–29^. While much research has focused on investigating the clinical relevance and pathogenesis of *E. coli* in the context of UTI, there is a paucity of studies investigating the pathogenesis of polymicrobial infection and opportunistic pathogens that frequently colonize catheterized patients. With the rise in antimicrobial resistance and the growing appreciation for the polymicrobial nature of CAUTI, there is a clear need for investigations into the impact of polymicrobial interactions as they may result in synergistic effects for co-colonizing pathogens ^30,31^.

Our prior work identified *P. mirabilis* and *E. faecalis* as the most common and persistent co-colonization partners in catheterized individuals ^10,24,32^, suggesting that interactions between these species facilitate persistent colonization. *P. mirabilis* is a Gram-negative rod-shaped bacterium that possess numerous virulence factors that contribute to the establishment of CAUTI and progression to secondary infections, and is the most common cause of infection-induced urinary stones, catheter encrustation, and blockage ^33–35^. *E. faecalis* is a Gram-positive, non-motile, and highly resistant bacterium of growing medical concern ^36–38^. Both species pose serious challenges to effective treatment that are compounded by co-colonization. It is therefore critical to understand the interactions between these two species and to identify potential strategies for disrupting persistent co-colonization.

We previously demonstrated that *P. mirabilis* and *E. faecalis* co-localize on catheters and within the bladder during experimental CAUTI, resulting in polymicrobial biofilms with enhanced biomass and antibiotic resistance ^10^. However, the underlying mechanism of biofilm enhancement was not elucidated. Coinfection with *P. mirabilis* and *E. faecalis* also dramatically increases the incidence of urolithiasis and bacteremia (39), although it is not yet known if the increase in disease severity is related to increased catheter biofilm biomass. In this study, we uncovered the metabolic interplay that drives biofilm enhancement and examined contribution to infection severity. We demonstrate that secretion of L-ornithine from *E. faecalis* via the ArcD arginine/ornithine antiporter drives L-arginine biosynthesis by *P. mirabilis,* ultimately increasing the protein content of polymicrobial biofilms and facilitating dissemination from the urinary tract to the bloodstream. Thus, modulating the metabolic interplay between these species could potentially disrupt polymicrobial biofilm formation, persistent colonization, and risk of progression to severe disease.

## Results

### *P. mirabilis* and *E. faecalis* polymicrobial biofilms have increased biofilm biomass that is associated with increased protein content

To investigate the underlying mechanism of enhanced biomass during polymicrobial biofilm formation, we began by studying single and polymicrobial biofilm formation in TSB-G (Tryptic soy broth supplemented with 1.5% glucose) under stationary conditions in 24-well plates. Biofilm biomass was assessed using crystal violet staining, while bacterial colony forming units (CFUs) were determined via serial dilution and plating on appropriate agar for each organism. *E. faecalis* forms slightly larger single-species biofilms than *P. mirabilis*; however, when grown together, biofilm biomass is significantly enhanced (Figure 1 A), confirming our previous observations ^39^. The increase in biofilm biomass was not driven by changes in total bacterial burden or viability as ∼10^8^ CFUs of each species were recovered from the single and co-culture biofilms (Figure 1 B). To determine the source of the increased biofilm biomass, we quantified the amount of protein, carbohydrate, and extracellular DNA (eDNA) in the total biofilm suspension (BS), the cell-associated fraction of the biofilm (CF), and the NaOH-extracted extrapolymeric substance fraction (EPS) from single and polymicrobial biofilms. Protein was the most abundant component of both the single-species and polymicrobial biofilms, and polymicrobial biofilms had a significant increase in total protein content in the biofilm suspension and cell-associated fraction compared to the single-species biofilms of *P. mirabilis* and *E. faecalis* (Figure 1 C). Carbohydrates were the next most abundant component of single-species and polymicrobial biofilms, but no significant increases were observed in the polymicrobial biofilms (Figure 1 D). The least abundant component of the biofilm matrix was found to be eDNA, and no significant increases in content were observed in the polymicrobial biofilms compared to the single-species (Figure 1 E). Thus, biofilm enhancement is driven by an increase in protein content stemming from the cell-associated fraction rather than the EPS. The importance of protein in mediating the enhancement phenotype was confirmed by establishing biofilms in the presence of 50 µg/mL of proteinase K (PK), which had no effect on single-species biofilms but resulted in a significant reduction in biomass of the polymicrobial biofilm (Figure 1 F).

**Figure 1.**
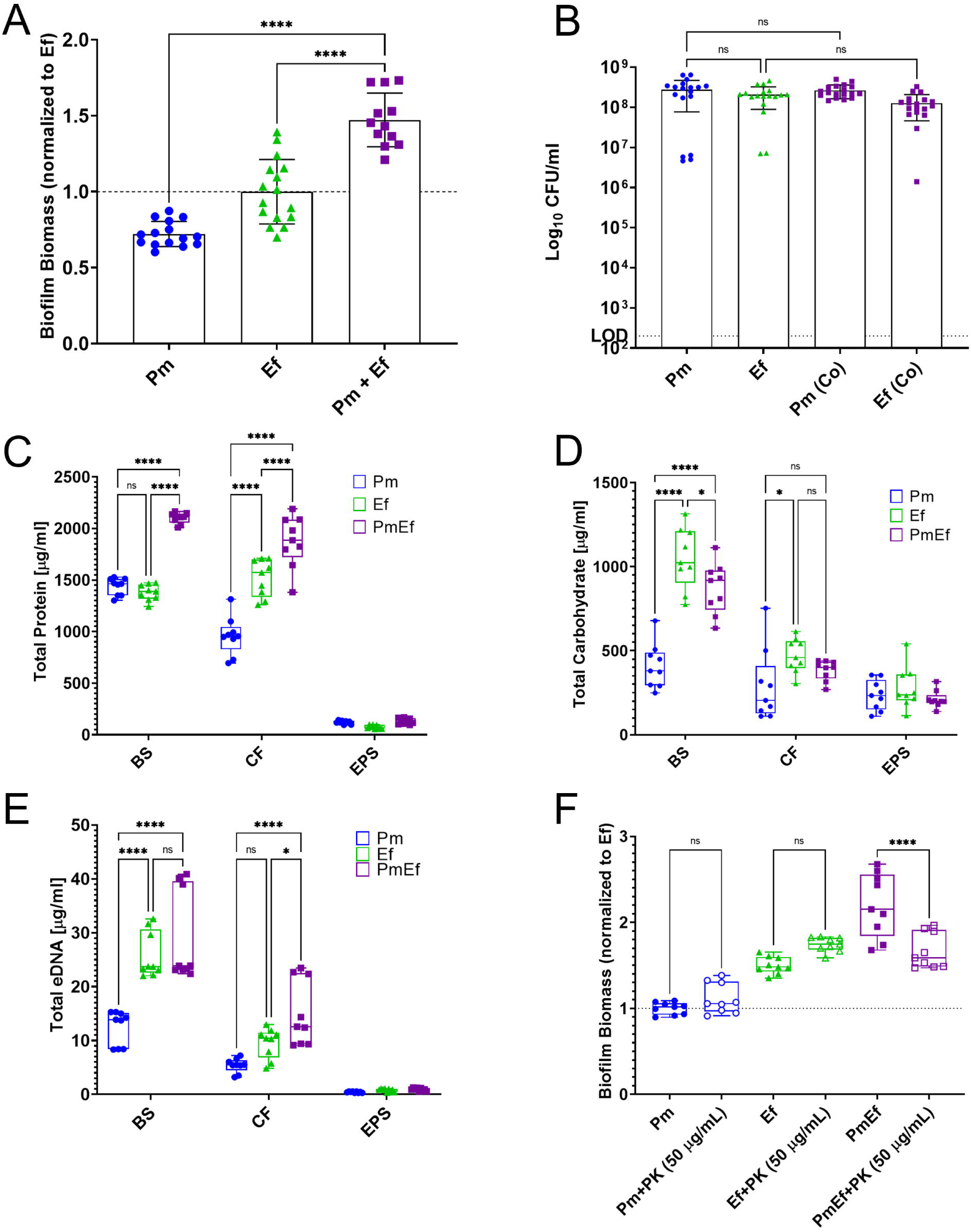
*P. mirabilis* and *E. faecalis* polymicrobial biofilms exhibit enhanced biofilm biomass that is driven by biofilm compositional changes and not bacterial counts. (A) Crystal violet staining of biofilms grown for 24hrs in TSB-G. (B) Colony forming units of biofilms grown for 24hrs in TSB-G. Data represent the mean ± standard deviation for 3-5 independent experiments with at least three replicates each. ns = non-significant, **** = P<.0001 by one-way ANOVA multiple comparisons. (C-E) Biofilm compositional analysis of 20-hour single or polymicrobial biofilms detailing C) total eDNA, D) total carbohydrate, and E) total protein, BS = biofilm supernatant fraction, CF = cell associated fraction, EPS = extrapolymeric substance fraction. (F) Crystal violet staining of biofilms grown for 24hrs in TSB-G with 50 µg/mL proteinase-K. Data represent the mean ± standard deviation for at least three independent experiments with at least two replicates each. ns = non-significant, * = P<.05; ** = P<.01; *** = P<.001; **** = P<.0001 by repeated measures one-way ANOVA.

Liquid chromatography mass spectrometry analysis of single and polymicrobial biofilms was next used to identify the proteins that are enriched in the polymicrobial biofilms. We identified 1427 proteins in the *P. mirabilis* single-species biofilms, 1061 proteins in *E. faecalis* single-species biofilms, and 1845 proteins in the polymicrobial biofilms, confirming an increase in protein content. Further, 78% of proteins from the polymicrobial biofilms mapped to *P. mirabilis* and 22% mapped to *E. faecalis*, suggesting that the majority of the biofilm protein content derives from *P. mirabilis.* This analysis revealed significant differences in protein abundances linked to a variety of metabolic pathways and virulence factors in *P. mirabilis* and *E. faecalis,* including an increase in abundance of multiple proteins related to ornithine and arginine biosynthesis and metabolism. (Table 1, Supplemental Table 1, and Supplemental Figure 1). Specifically, 37 *P. mirabilis* proteins were enriched greater than 2-fold in the polymicrobial biofilm compared to *P. mirabilis* single biofilms, and 6/25 are involved in ornithine/arginine transport and metabolism. In *E. faecalis,* 225 proteins were enriched (>2-fold) compared to the single biofilm, the majority of which pertain to metabolism, translation, and cell growth and division. The changes in ornithine and arginine metabolism drew immediate interest in light of previous work by Keogh et al, wherein L-ornithine export from *E. faecalis*, driven by the ArcD L-ornithine/L-arginine antiporter, modulated biofilm formation, siderophore production, and fitness of *E. coli* ^40^. *P. mirabilis* can either directly metabolize L-ornithine to the polyamine putrescine via ornithine decarboxylase (SpeF), or it can use L-ornithine for L-arginine biosynthesis via ornithine carbamoyltransfers (ArgI/ArgF), and can then catabolize L-arginine to putrescine via arginine decarboxylase (SpeA) and agmatinase (SpeB). We therefore focused on the contribution of this pathway to polymicrobial biofilm enhancement.

**Table 1.**
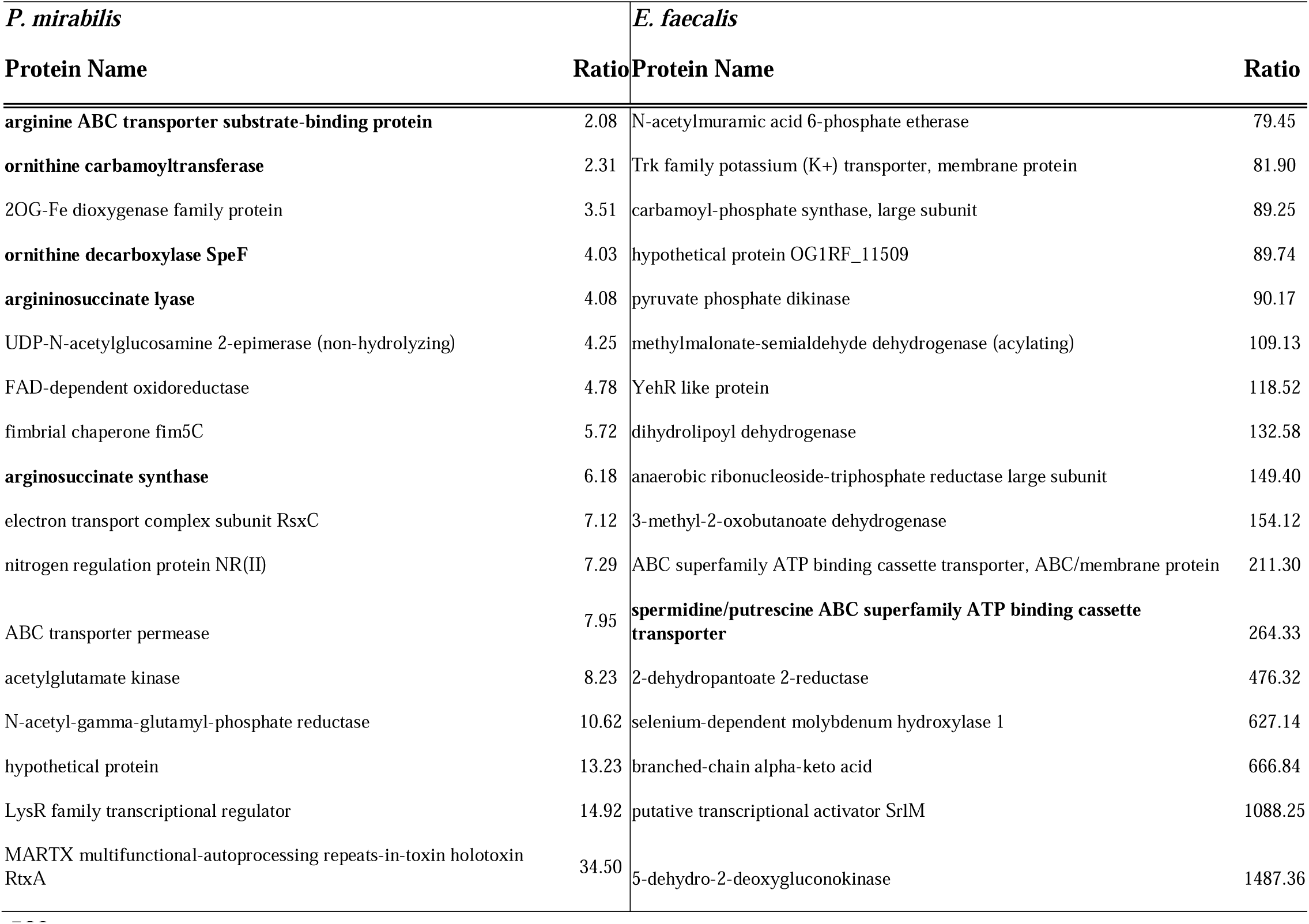
Liquid chromatography mass spectrometry (LC-MS) analysis shows an upregulation of many ornithine and arginine metabolism related genes. LC-MS analysis summary detailing the ratio of select proteins within the polymicrobial biofilms to single species biofilms. Select proteins that were at least 2-fold increased in polymicrobial biofilms are detailed within, proteins related to ornithine/arginine metabolism are bolded.

### Arginine biosynthesis from ornithine is a critical determinant of *P. mirabilis* fitness *in vitro*

Before investigating the importance of arginine/ornithine metabolism in mediating the biofilm enhancement phenotype, we first examined the growth characteristics of an *E. faecalis arcD* mutant as well as *P. mirabilis* ornithine catabolism mutants *speF* (PMI0307) and *argI/F* (PMI3457, herein referred to as *argF*) under relevant conditions. Disrupting ornithine export had no impact on *E. faecalis* growth or viability as the *arcD* mutant grew similarly to wild-type OG1RF in brain heart infusion broth (BHI), tryptic soy broth supplemented with 1.5% glucose (TSB-G), and pooled human urine (Figure 2 A, B, & C). Similarly, disrupting ornithine metabolism and arginine biosynthesis by *P. mirabilis* had no impact on growth or viability in rich media, as the *argF* and *speF* mutants grew similarly to wild-type HI4320 in TSB-G and Luria-Bertani broth (LB) (Figure 2 D & E). However, in minimal salts media (PMSM), loss of *argF* completely abrogated *P. mirabilis* growth while loss of *speF* had no impact (Figure 2 F). The *argF* mutant growth defect could be fully rescued by supplementation with either L-citrulline or L-arginine but not by L-ornithine or the arginine catabolic products agmatine or putrescine, demonstrating that mutation of *argF* results in L-arginine auxotrophy in *P. mirabilis* (Figure 2 F).

**Figure 2.**
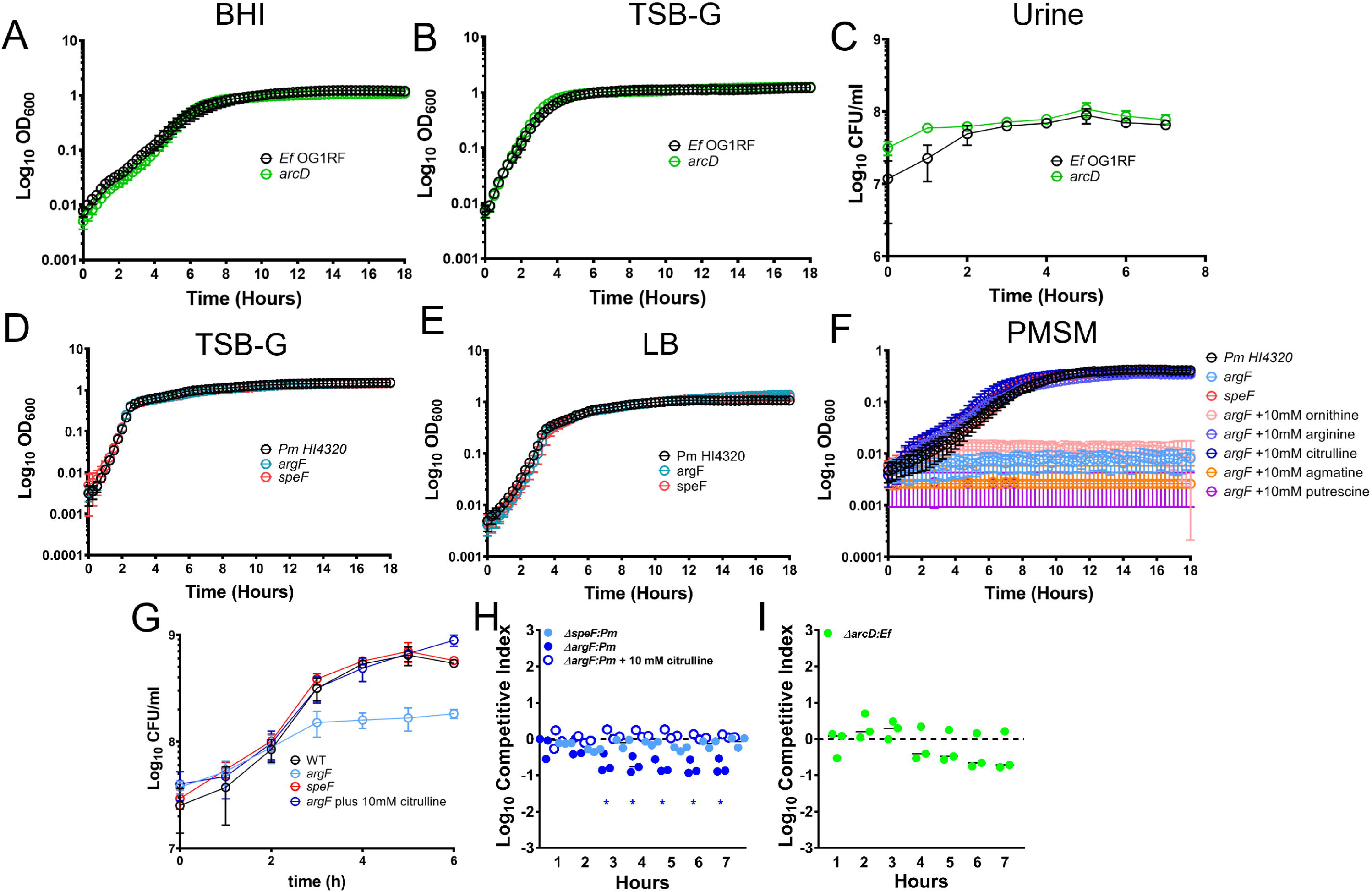
Ornithine and arginine metabolism are key determinants of *P. mirabilis* growth in vitro in both laboratory media and physiologically relevant pooled human urine. *E. faecalis* growth curves in A) BHI, B) TSB-G, and C) pooled human urine. *P. mirabilis* growth curves in D) TSB-G E) LB, F) PMSM (*Proteus* minimal salts media) without supplementation or supplemented with 10 mM ornithine, 10mM arginine, 10 mM citrulline, 10 mM agmatine, or 10 mM putrescine. G) CFUs of *P. mirabilis* and mutants during growth in pooled human urine. (H-I) Competitive index (CI) of *P. mirabilis* mutants co-inoculated with wild-type *P. mirabilis* (H) or *E. faecalis arcD* mutant co-inoculated with wild-type *E. faecalis* (I) in pooled human urine. Each symbol represents the log10 CI for an individual inoculum, error bars represent the medians and dashed line indicates the log10 CI = 0 (the expected value if the ratio of mutant/WT is 1:1). * = P<.05 by one sample t test against a theoretical log10 CI = 0.

Human urine is considered to be a nutrient-limited medium for bacteria, providing mainly amino acids and small peptides as nutrient sources ^41^. When grown in pooled human urine, the *speF* mutant grew similarly to wild-type while viability of the *argF* mutant stopped increasing after ∼3 hours (Figure 2 G). This corresponds to the timing of a ∼50% reduction in the arginine concentration of urine (from ∼134 µM to ∼60 µM) by wild-type *P. mirabilis* ^42^, which is notable as a prior study observed that growth of an *E. coli* arginine auxotroph became limited when the concentration of arginine decreased below 60 µM ^43,44^. Importantly, growth of *argF* in human urine could again be rescued by supplementation with L-citrulline to fuel L-arginine biosynthesis, much like growth in minimal medium (Figure 2 G).

Subtle fitness defects can often be further magnified when a mutant strain is directly competing against its parental isolate. We therefore conducted co-challenge experiments in human urine, in which cultures were inoculated with a 1:1 mixture of each mutant versus wild-type and a competitive index was calculated based on their ratio at the start of the experiment and hourly thereafter (Figure 2 H & I). During direct co-challenge with wild-type *P. mirabilis,* fitness of the *argF* mutant was significantly decreased after just 2 hours of growth in urine. The defect was likely due to competition for arginine and was rescued by supplementation with citrulline. In contrast, no fitness defects were observed for the *speF* mutant or the *E. faecalis arcD* mutant. Thus, the use of L-ornithine to fuel L-arginine biosynthesis is critical for *P. mirabilis* growth in minimal medium and for optimal fitness in human urine, but ornithine secretion by *E. faecalis* and ornithine catabolism to putrescine in *P. mirabilis* are dispensable.

We previously demonstrated that *P. mirabilis* and *E. faecalis* do not exhibit obvious competitive behavior in human urine, as the growth rates for each species were equivalent during co-culture compared to single-species culture ^45^. However, considering the importance of L-arginine biosynthesis to *P. mirabilis* fitness during growth in urine, we sought to determine whether arginine/ornithine interplay alters viability of either species during co-culture in urine (Supplemental Figure 2 A). Interestingly, growth of the *argF* mutant plateaued early during co-culture wild-type *E. faecalis* but not during co-culture with the *arcD* mutant. This observation suggests that *E. faecalis* may impair growth of the *argF* mutant by stealing the limited L-arginine present in urine. In contrast, the *E. faecalis arcD* mutant exhibited a slightly faster initial growth rate during co-culture with *P. mirabilis* than wild-type *E. faecalis,* and this was independent of *P. mirabilis* L-arginine biosynthesis (Supplemental Figure 2 B). Thus, loss of *arcD* may provide *E. faecalis* with a slight advantage during initial co-culture with *P. mirabilis,* although the mutant and wild-type strains achieved the same final cell density in stationary phase.

### L-ornithine secretion by *Ef* facilitates polymicrobial biofilm enhancement

To investigate the contribution of *Ef* arginine/ornithine antiport to polymicrobial biofilm enhancement, we established single and polymicrobial biofilms with *Pm,* Δ*argF, Ef,* and Δ*arcD* and measured biofilm biomass and protein content. Neither of the mutants exhibited differences in single-species biofilm biomass compared to their respective parental strains (Figure 3C). However, enhancement of biofilm biomass and protein content were both abrogated during co-culture of *Pm* with Δ*arcD* (Figure 3C and D), indicating that arginine/ornithine antiport by *Ef* is critical for the increased biomass that occurs during co-culture. When *Pm* Δ*argF* was co-cultured with wild-type *Ef,* biofilm enhancement was still observed but to a lower level than for the parental strains, and protein levels were similar. Thus, the ability of *Pm* to use L-ornithine for production of citrulline during L-arginine biosynthesis is not critical for biofilm enhancement under these conditions. Importantly, all differences in biofilm biomass and protein content were independent of any potential impact on bacterial viability (Supplemental Figure 3A).

**Figure 3.**
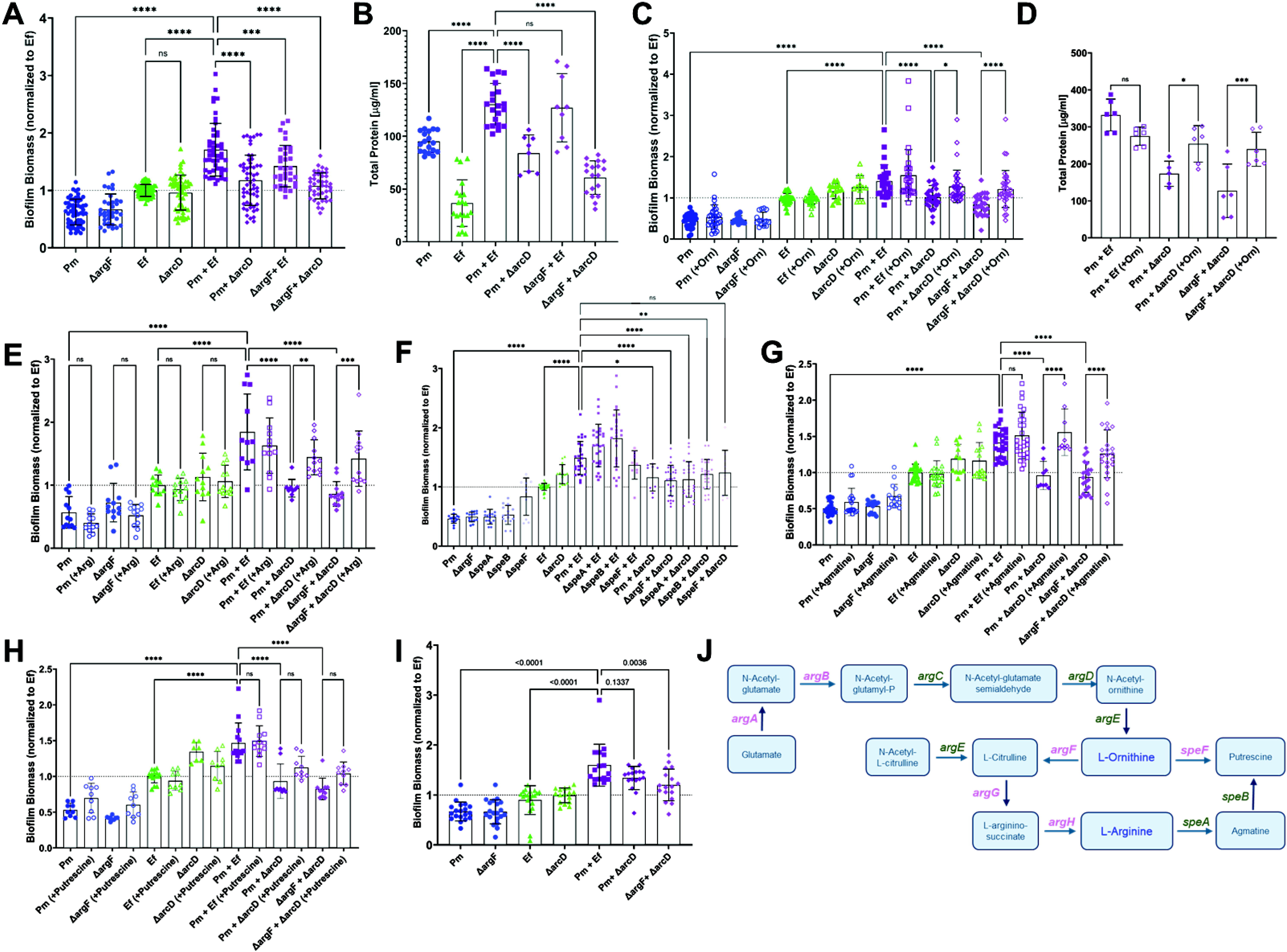
L-ornithine secretion from E*. faecalis* drives *P. mirabilis* arginine biosynthesis and increased biofilm biomass. (A) Crystal violet staining of single-species and polymicrobial biofilms grown for 24 hrs in TSB-G. (B) Total protein content as measured by BCA from three pooled biofilms per experiment. (D) Crystal violet staining of biofilms grown for 24 hrs in TSB-G with or without 10 mM of L-ornithine. (D) Total protein content as measured by BCA from three pooled biofilms per experiment when established in TSB-G with or without 10mM L-ornithine supplementation. (E) Crystal violet staining of biofilms grown for 24 hrs in TSB-G with or without 10 mM of L-arginine. (F) Crystal violet staining of biofilms grown for 24 hrs in TSB-G with *P. mirabilis* arginine catabolism mutants, *speA* and *speB*. (G-H) Crystal violet staining of biofilms grown for 24 hrs in TSB-G with or without 10 mM of agmatine (G) or putrescine (H). (I) Crystal violet staining of single or polymicrobial biofilms grown for 24 hours in pooled human urine. Data represent the mean ± SD for 3-5 independent experiments with at least two replicates each. ns = non-significant, One-way ANOVA multiple comparisons, * p < 0.05, ** p < 0.01, *** p < 0.001, **** p <0.0001. (J) All known genes involved in L-ornithine metabolism and L-arginine biosynthesis in *P. mirabilis* are displayed. L-ornithine can either be directly catabolized to putrescine via ornithine decarboxylate (SpeF), or it can feed into L-arginine biosynthesis via ornithine carbamoyltransferse (ArgF), which generates L-citrulline. Argininosuccinate synthase (ArgG) uses ATP to generate L-arginino-succinate from L-citrulline and L-aspartate, then argininosuccinate lyase (ArgH) generates L-arginine and fumarate from L-arginino-succinate. L-arginine can then be catabolized to putrescine via arginine decarboxylase (SpeA) and agmatinase (SpeB). Genes in purple were identified as enriched in polymicrobial biofilms.

We previously demonstrated that direct cell-cell contact was required for biofilm enhancement, as neither *Pm* nor *Ef* exhibited altered biofilm biomass during co-culture when separated by a transwell insert ^39^. Thus, it was surprising that loss of arginine/ornithine antiport in *Ef* abrogated polymicrobial biofilm enhancement. We therefore sought to determine if exogenous ornithine could promote biofilm enhancement. In agreement with our prior findings, the addition of 10 mM ornithine had no impact on single species biofilm biomass for any of the strains (Figure 3E). However, ornithine supplementation fully restored biofilm enhancement during co-culture of either wild-type *Pm* or Δ*argF* with *Ef* Δ*arcD* (Figure 3E) and also restored biofilm protein levels (Figure 3F). Thus, the presence of excess ornithine alone is sufficient to restore contact-dependent enhancement of biofilm biomass during co-culture. If ornithine-dependent arginine biosynthesis in *Pm* was required for biofilm enhancement, supplementation should not have restored enhancement during co-culture of Δ*argF* with Δ*arcD.* Considering that auxotrophy of the Δ*argF* mutant could not be complemented by supplementation with ornithine, these findings suggest that ornithine either promotes biofilm enhancement through a mechanism that is independent of *Pm* arginine biosynthesis, or that *Pm* has access to alternative precursors for arginine biosynthesis during co-culture with *Ef*.

To examine the specific contribution of arginine to biofilm enhancement, supplementation experiments were repeated with 10 mM L-arginine (Figure 3G). Supplementation again had no impact on single species biofilms, but the addition of arginine restored biofilm enhancement during co-culture of either wild-type *Pm* or Δ*argF* with *Ef* Δ*arcD.* Considering that *Ef* encodes other arginine import systems such as the Art ABC transporter, excess arginine could still be taken up by *Ef* without ornithine antiport. Thus, arginine import by either *Pm* or *Ef* restores contact-dependent biofilm enhancement.

To determine if arginine catabolism or putrescine biosynthesis by *Pm* are required for biofilm enhancement, we next used *Pm* mutants in *speA*, *speB*, and *speF* (Figure 3H) ^46^. Much like Δ*argF*, single species biofilms formed by each of the mutants exhibited similar biomass to wild-type *Pm*. Polymicrobial biofilms formed with each of the mutants were identical to those formed by wild-type *Pm,* indicating that L-arginine catabolism and putrescine biosynthesis are not required for polymicrobial biofilm enhancement. To confirm these results, we further examined the contribution of agmatine and putrescine to polymicrobial biofilm enhancement, as *Ef* produces an agmatine/putrescine antiporter. Supplementation with 10 mM agmatine had no impact on single species biofilms, but fully restored contact-dependent biofilm enhancement during co-culture of either wild-type *Pm* or Δ*argF* with *Ef* Δ*arcD* (Figure 3I). In contrast, supplementation with putrescine failed to restore biofilm biomass (Figure 3J). Since *Pm* can only use agmatine to produce putrescine and neither putrescine supplementation nor loss of agmatinase activity (Δ*speB*) abrogated enhancement, our findings suggest that agmatine is most likely mediating enhancement via import by *Ef.* Taken together, these data suggest that biofilm enhancement is mediated through a combination of ornithine production by *Ef,* agmatine import by *Ef,* and arginine import by at least one species, all of which are disrupted by loss of arginine/ornithine antiport in *Ef*.

### L-ornithine secretion by *E. faecalis* and L-arginine biosynthesis by *P. mirabilis* contribute to enhanced disease severity and dissemination during polymicrobial infection

We previously demonstrated that polymicrobial infection with *E. faecalis* and *P. mirabilis* increases disease severity during experimental CAUTI and we demonstrated the importance of biofilm formation to bacterial pathogenesis in the context of CAUTI ^39,45,46^. We therefore sought to determine the contribution of *E. faecalis* L-ornithine secretion and *P. mirabilis* L-arginine biosynthesis to establishing polymicrobial catheter biofilms as well as promoting dissemination to the kidneys and bloodstream and overall disease severity. We utilized the *E. faecalis arcD* mutant and the *P. mirabilis argF* mutant to examine the specific contribution of ornithine export and arginine biosynthesis to pathogenesis in the well-established murine CAUTI model ^42,47,48^. Female CBA/J mice aged 6-8 weeks were transurethrally inoculated with 10^5^ CFUs of either wild type *P. mirabilis*, the *argF* mutant, wild-type *E. faecalis,* the *arcD* mutant, or polymicrobial mixtures, and a 4mm silicone catheter segment was placed in the bladder during inoculation. Mice were euthanized 96 hours post-inoculation and bacterial burden was quantified in the urine, bladder, kidneys, and spleen (Figure 4).

**Figure 4.**
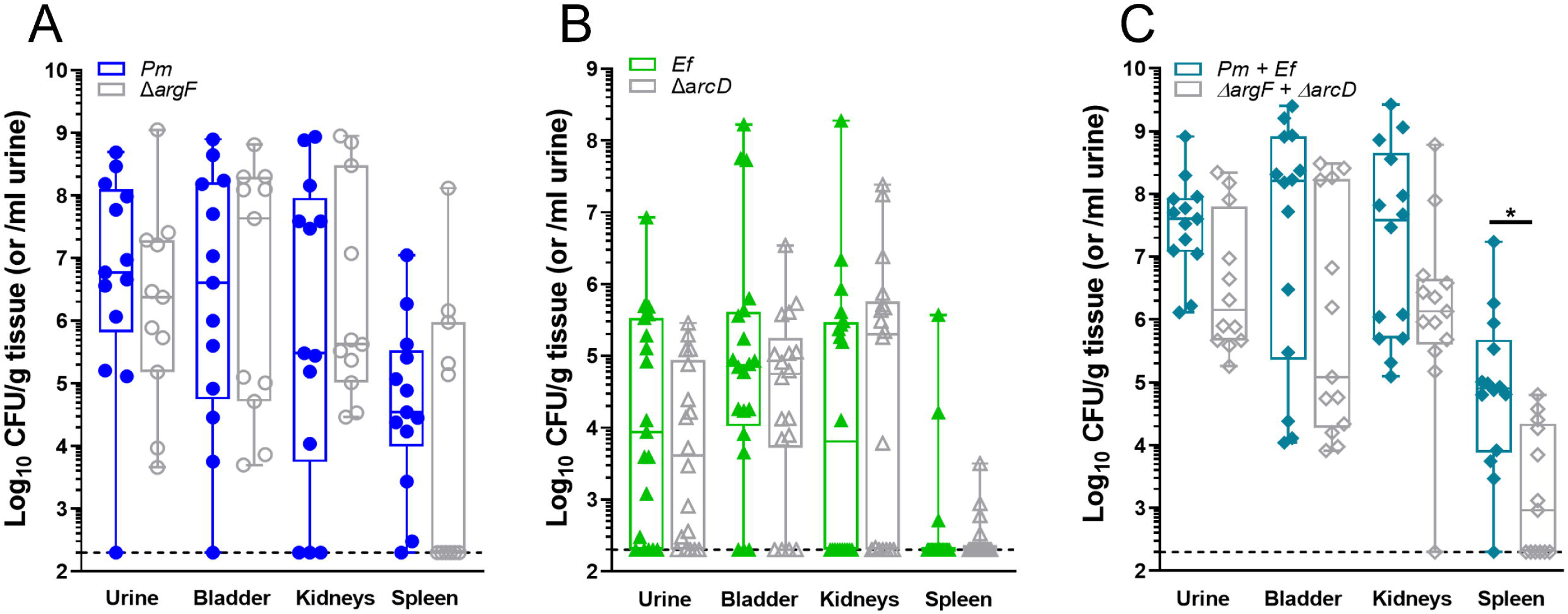
Metabolic interplay between *E. faecalis* and *P. mirabilis* contributes to secondary bacteremia. Bacterial counts in urine, bladder, kidney, and spleen samples collected at 96hrs post infection in a CAUTI murine model. Animals were infected with 10^5^ CFUs/mL of either A) wild type *P. mirabilis* or *P. mirabilis* Δ*argF*, or B) wild type *E. faecalis* or *E. faecalis* Δ*arcD* in single species infections. (C) Mice were coinfected with a 50:50 mixture of the wild-type strains or their respective mutants for polymicrobial infection experiments, with bacterial counts being depicted as total bacterial burden per organ. Total bacterial burden was analyzed via nonparametric One-Way ANOVA, * p = 0.0154. Data presented is representative of three combined, independent animal studies, n = 4-16.

Neither ArgF nor ArcD alone were important for establishing single-species infection, as the mutants colonized all organs of the urinary tract to a similar level as the wild-type strains (Figure 4 A & B). There were also no differences in infection severity for *arcD* compared to wild-type *E. faecalis,* although there was a decrease in the number of mice that developed bacteremia during infection with the *argF* mutant compared to wild-type *P. mirabilis* (Table 2). Differences in disease severity became more apparent in the context of polymicrobial infection. While both species were detected in all coinfected mice by differential plating (Supplemental Figure 4), mice coinfected with the mutant strains displayed a trend towards decreased CFUs in all organs compared to mice coinfected with the wild-type strains, which was statistically significant in the spleen (Figure 4 C). Strikingly, animals coinfected with the mutant strains had significantly fewer indicators of severe disease as compared to mice coinfected with the wild type strains, including kidney discoloration and mottling, kidney hematoma, and bacteremia (Table 2). The decreased incidence of bacteremia is particularly notable, as an increased incidence of bacteremia is one of the hallmarks of *P. mirabilis* and *E. faecalis* coinfection compared to single-species infection ^45^. Taken together, these data clearly demonstrate that L-ornithine secretion by *E. faecalis* facilitates L-arginine biosynthesis by *P. mirabilis,* the combined action of which alters polymicrobial biofilm formation and infection severity.

**Table 2.** Metabolic interplay between *E. faecalis* and *P. mirabilis* contributes to polymicrobial infection severity. Incidence of bacteremia and tissue abnormalities in murine CAUTI infections. Chi-square tests were used to determine if frequency of events are statistically different between groups.

| Adverse health event or tissue abnormality | *Pm* | $\Delta argF$ | Chi-square<br>P value | *Ef* | $\Delta arcD$ | Chi-square<br>P value | *Pm* +<br>*Ef* | $\Delta argF$ +<br>$\Delta arcD$ | Chi-square<br>P value |
| --- | --- | --- | --- | --- | --- | --- | --- | --- | --- |
| Bladder hematoma and/or blood in urine | 1/13 | 0/11 | 0.306 | 0/20 | 0/18 | >0.999 | 0/14 | 0/13 | >0.999 |
| Kidney hematoma | 3/13 | 0/11 | 0.088 | 0/20 | 0/18 | >0.999 | 5/14 | 0/13 | 0.011* |
| Kidney color change and/or mottling | 1/13 | 0/11 | 0.347 | 0/20 | 0/18 | >0.999 | 8/14 | 2/13 | 0.013* |
| Kidney stone | 2/13 | 3/11 | 0.475 | 0/20 | 0/18 | >0.999 | 5/14 | 1/13 | 0.054 |
| Any abnormality | 4/13 | 3/11 | 0.099 | 0/20 | 0/18 | >0.999 | 8/14 | 2/13 | 0.013* |
| Bacteremia | 11/13 | 4/11 | 0.015* | 0/20 | 0/18 | >0.999 | 13/14 | 6/13 | 0.003* |

## Discussion

Bacterial biofilms have long been noted to be vital for pathogenesis and disease progression in a variety of disease contexts, including CAUTI ^49–52^. There has been a growing appreciation for the fact that many diseases and biofilms are polymicrobial environments, where the network of interactions between bacterial species and the host are important determinants of the overall course of disease development ^10,32,52–55^. However, there is still a paucity of studies addressing the interactions that contribute to polymicrobial biofilm formation, colonization, and pathogenesis. Previously, we demonstrated that *E. faecalis* and *P. mirabilis* are frequent co-colonizers in catheterized patient populations and that they co-localize and form unique biofilm communities with enhanced biofilm biomass, persistence, and antibiotic resistance ^10^. However, the underlying mechanism was yet to be understood.

Herein, we have demonstrated that L-ornithine secretion from *E. faecalis* feeds into L-arginine biosynthesis and metabolism by *P. mirabilis,* resulting in a contact-dependent increase in protein content and biofilm biomass compared to single-species biofilms (summarized in Figure 5). Not only were we able to demonstrate that this metabolic interaction influences biofilm formation *in vitro*, but we also uncovered an important role for this metabolic interplay in mediating disease severity in a mouse model of polymicrobial CAUTI. This study adds to a growing body of work that metabolic cross-feeding is a determinant of polymicrobial infections. Previously, Keogh et al (2016) demonstrated that L-ornithine secretion from *E. faecalis* increased *E. coli* siderophore production and biofilm growth under iron-limitation, as well as persistence in a wound infection model ^40^. More recently, work by Smith et al. (2022) demonstrated that *E. faecalis* enhances the fitness and virulence of the gut pathogen *Clostridioides difficile* by providing a source of fermentable amino acids, including ornithine ^56^. In both of these studies, L-ornithine from *E. faecalis* enhanced growth of the partner species. In contrast, our work demonstrates a growth-independent role for L-ornithine in mediating contact-dependent polymicrobial interactions. Together, these studies underscore the pivotal role that *E. faecalis* L-ornithine secretion plays in mediating different polymicrobial interactions in multiple disease contexts, and further highlights the potential of L-ornithine as a common metabolite cue.

**Figure 5.**
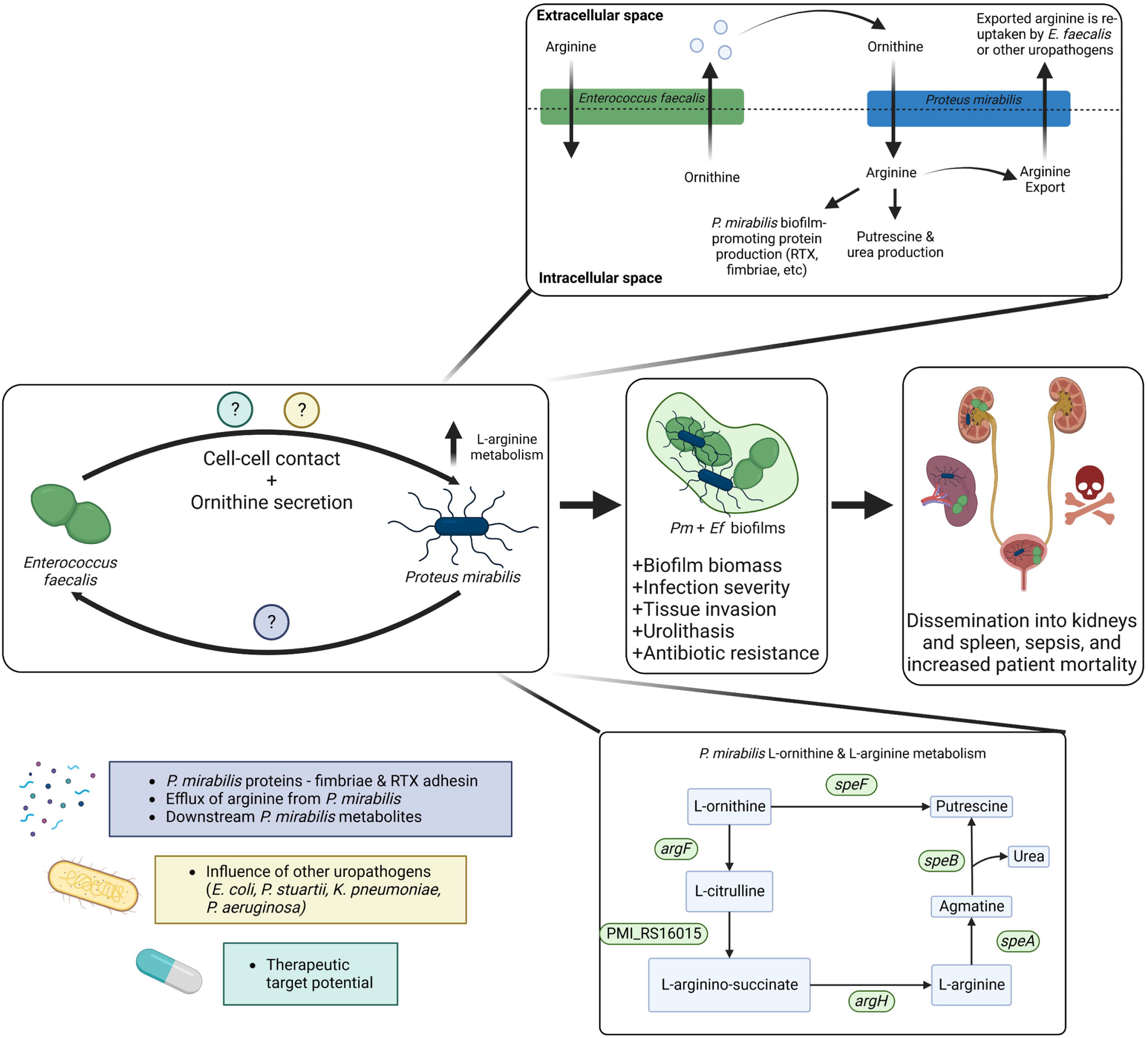
Working model of the metabolic interplay between *E. faecalis* and *P. mirabilis* in polymicrobial CAUTI infection. Model of metabolic interplay between *E. faecalis* and *P. mirabilis* in polymicrobial biofilms. All known genes involved in L-ornithine metabolism and L-arginine biosynthesis in *P. mirabilis* are displayed. L-ornithine can either be directly catabolized to putrescine via ornithine decarboxylate (SpeF), or it can feed into L-arginine biosynthesis via ornithine carbamoyltransferse (ArgF), which generates L-citrulline. Argininosuccinate synthase (PMI_RS16015) uses ATP to generate L-arginino-succinate from L-citrulline and L-aspartate, then argininosuccinate lyase (ArgH) generates L-arginine and fumarate from L-arginino-succinate. L-arginine can then be catabolized to putrescine via arginine decarboxylase (SpeA) and agmatinase (SpeB). Our data support a model in which ornithine secretion by *E. faecalis* coupled with direct cell-cell contact increases L-arginine metabolism and protein expression by *P. mirabilis*, leading to the development of a polymicrobial biofilm with significantly increased biomass and antibiotic resistance. Co-colonization of the two pathogens increases morbidity and mortality in a murine CAUTI model. However, disruption of ornithine/arginine metabolic interplay leads to significant reductions disease severity, revealing a new potential target for disrupting polymicrobial infection.

Considering that supplementation with either ornithine, arginine, or agmatine restored polymicrobial biofilm enhancement for the mutant strains while putrescine did not, our data indicate that either intracellular L-arginine stores or biosynthesis intermediates are likely the key mediators of biofilm enhancement in *Pm*. Our prior studies have demonstrated an important contribution of L-arginine to *P. mirabilis* fitness and virulence; specifically, L-arginine acts as an environmental cue to promote swarming motility, and catabolism to agmatine via SpeA contributes to acid tolerance, motility, and fitness within the urinary tract in addition to fueling putrescine biosynthesis ^57^. In our prior genome-wide transposon insertion site sequencing (Tn-seq) study, we also found that polymicrobial infection with another common coinfection partner, *Providencia stuartii,* causes *P. mirabilis* to require the L-arginine biosynthetic pathway (including *argF*) but not *speA, speB,* or *speF* for optimal fitness, suggesting a specific involvement of L-arginine and its biosynthesis intermediates rather than its catabolic products. ^58^. Combined with this current study, these observations hint at the potential for L-arginine to act as a key determinant of *P. mirabilis* virulence and fitness in polymicrobial CAUTI.

The observation that ornithine supplementation restored enhancement of biofilms formed by co-culture of Δ*argF* with Δ*arcD* was initially surprising since disrupting ornithine carbamoyltransferase should prevent generation of citrulline and subsequently arginine by *Pm*. However, it is possible that ornithine catabolism via *speF* may at least partially compensate for loss of *argF* under these conditions. Unfortunately, testing this hypothesis requires an *argF/speF* double mutant in *Pm* for which numerous attempts proved unsuccessful, suggesting that ornithine catabolism by at least one of these pathways is required for *Pm* viability *in vitro*.

While the specific fate of arginine remains to be determined, we hypothesize that *P. mirabilis* may directly utilize the excess L-arginine generated from L-ornithine for production of specific proteins that mediate biofilm enhancement. Our proteomics experiments revealed a 5-fold increase in the fimbrial chaperone protein fim5C in polymicrobial biofilms compared to *P. mirabilis* single-species biofilms, suggesting that L-arginine may contribute to production of certain fimbriae (Table 1). Fimbriae are known to play a vital role in *P. mirabilis* biofilm formation and mediate adherence to the catheter surface ^33,59^, although their specific contribution to polymicrobial biofilm formation has yet to be explored. Another product of *P. mirabilis* that may be responsible for the increase in biofilm biomass in the polymicrobial biofilms is a putative repeats-in-toxin (RTX) adhesion protein RtxA. RTX toxins are part of a family of pore forming cytolysins produced mainly by Gram-negative bacteria, and they can have diverse functions including adhesion, which can play a role in biofilm formation ^60–62^. The most well characterized RTX adhesin is the *Pseudomonas fluorescens* protein LapA, which was shown to be essential for biofilm formation ^62,63^. RtxA was the most over-represented *P. mirabilis* protein from polymicrobial biofilms, suggesting that *E. faecalis* enhances RtxA production during biofilm formation. Interestingly, both *fim5C* and *rtxA* were identified as *P. mirabilis* fitness factors during polymicrobial CAUTI with *P. stuartii* but not during single-species infection, suggesting that both proteins may play a specific role in mediating polymicrobial interactions ^58^. The contribution of these adhesins to polymicrobial interactions are an active area of ongoing investigation.

As an alternative, the L-arginine biosynthesis intermediates generated from L-ornithine may be acting as nutrient signals within *P. mirabilis* and triggering phenotypic changes that ultimately influence biofilm formation, rather than being directly used for protein synthesis. It is also possible that the L-arginine generated by *P. mirabilis* is being exported and taken up by *E. faecalis* through an *arcD*-independent transport mechanism, and that *E. faecalis* is driving the contact-dependent biofilm enhancement through an arginine-dependent mechanism. The full complement of arginine import and export machinery are not yet fully elucidated in these species, but uncovering the mechanisms behind arginine export in *P. mirabilis* and import by *E. faecalis* are expected to contribute to a deeper understanding into this important polymicrobial interaction. Future efforts will be focused on identifying the specific protein mediators of biofilm enhancement, distinguishing which microbe is responsible for their production, and defining the role of L-arginine in their biosynthesis.

It is also important to remember that the CAUTI bladder environment is typically polymicrobial and thus there are many more bacterial species present than just *P. mirabilis* and *E. faecalis* ^56,64–66^. Bacteria engage in multiple cooperative and competitive interactions, which can be mediated by small molecules such as L-ornithine, and multiple different bacterial species can be competing for and responding to the same metabolic cue. The addition of other uropathogens is likely to modulate or influence the cross-feeding interaction described here, and this can be accomplished through a variety of mechanisms that change the nutrient landscape, spatial structure of the community, or community metabolism ^67,68^. Additionally, other uropathogens, such as *P. stuartii, E. coli,* and *Morganella morganii,* have been shown to modulate virulence factor production and activity in *P. mirabilis* and may also contribute to biofilm formation ^45,47,69^. It is also possible that with the addition of other common uropathogens, the L-ornithine-driven biofilm enhancement may be disrupted though competition for this or other metabolites.

Given that *P. mirabilis* and *E. faecalis* both exhibit intrinsic resistance to several antibiotics and that polymicrobial biofilm formation further exacerbates these concerns, this work can be used to develop new approaches to prevent or disrupt biofilm formation. Our data suggest that targeting ornithine metabolism and arginine biosynthesis may represent a new avenue for exploration, especially as disrupting these pathways also decreases risk of developing severe disease during experimental infection. However, considering that catheter-associated bacteriuria and CAUTI also frequently involve additional co-colonizing species such as *E. coli,* additional investigations are needed to understand how other co-colonizing pathogens influence biofilm formation, metabolic cross-feeding, and disease progression.

## Methods

### Bacterial strains

*Proteus mirabilis* strain HI4320 was isolated from the urine of a long-term catheterized patient in a chronic care facility ^70^. All *P. mirabilis* mutants used in this study were generated by inserting a kanamycin resistance cassette into the gene of interest following the Sigma TargeTron group II intron protocol as previously described ^71,72^. The *P. mirabilis speF, speA,* and *speB* mutants were previously constructed ^57^, while the *argF* mutant was specifically generated for this study. Mutants were verified by selection on kanamycin and PCR. The *Enterococcus faecalis* strain used in this study is an oral clinical isolate, OG1RF ^73,74^. The *E. faecalis* Δ*arcD* mutant was previously generated via mariner transposon mutagenesis ^40,75,76^.

### Bacterial culture conditions

*P. mirabilis* was cultured at 37°C with shaking at 225 RPM in 5 mL of low-salt LB (LSLB) broth (10 g/L tryptone, 5 g/L yeast extract, 0.1g/L NaCl) or on LSLB plates solidified with 1.5% agar. *E. faecalis* was routinely cultured in 5 mL of Brain Heart Infusion (BHI) broth at 37°C with shaking at 225 RPM or on BHI agar plates solidified with 1.5% agar. *P. mirabilis* mutant strains were grown in media supplemented with 50 µg/mL kanamycin, while the *E. faecalis* mutant strain was grown with 8 µg/mL chloramphenicol. Both species of bacteria were also grown in Tryptic Soy Broth supplemented with 1.5% glucose (TSB-G) as indicated. *Proteus mirabilis* minimal salts medium (PMSM) was used in experiments requiring defined growth medium (10.5 g/L K_2_HPO_4_, 4.5 g/L KH_2_PO_4_, 1 g/L (NH_4_)_2_SO_4_, 15 g/L agar, supplemented with 0.002% nicotinic acid, 1 mM MgSO_4_, and 0.2% glycerol). PMSM was supplemented further with addition of 10 mM L-ornithine, L-arginine, or L-citrulline as indicated. Filter sterilized pooled human urine from at least 20 de-identified female donors was purchased from Cone Bioproducts (Sequin, TX), stored at -20C, and used as indicated as a physiologically relevant growth medium. For co-challenge experiments, samples were plated on plain LSLB agar (total CFUs), LSLB with kanamycin (*P. mirabilis* mutant strain CFUs), and BHI agar supplemented with 100 µg/ml spectinomycin (*E. faecalis* CFUs).

### Crystal violet staining of bacterial biofilms

Overnight cultures of wild-type or mutant bacteria were adjusted to approximately 2x10^7^ CFU/mL, an OD_600_ of 0.02 for *P. mirabilis* and OD_600_ of 0.04 for *E. faecalis*, in either TSB-G or pooled human urine as indicated, and 750 µL was dispensed in triplicate into the wells of tissue culture treated 24-well plates (Falcon 353047). For polymicrobial biofilms, 325 µL of the appropriate *P. mirabilis* and *E. faecalis* strains were added to the well to a final volume of 750 µL. Sterile media was dispensed in triplicate into wells to serve as a blank for the crystal violet staining. Plates were incubated for 24 hours at 37°C in partially sealed bags with a damp paper towel, after which supernatants were gently aspirated and adherent biofilms were washed twice with 1 mL of 1x phosphate buffered saline (PBS), with care taken to not disrupt the biofilm community. Next, 1 mL of 95% ethanol was added to each well and the plate was incubated at room temperature for 15 minutes, after which ethanol was aspirated and the plate was allowed to air dry with the lid off for 60 minutes. Biofilms were then stained with 0.1% crystal violet and incubated at room temperature for 60 minutes, after such time the crystal violet solution was aspirated and biofilms were washed once with 1 mL of deionized water. Stained biofilms were solubilized in 1 mL of 95% ethanol and plates were incubated at room temperature on a plate shaker at 200 RPM for 15 minutes. Using a 1 mL micropipette tip, the bottom and sides of wells in the plate were scraped to ensure all stained biofilm biomass was fully resuspended. Crystal violet absorbance was then read at 570 nm using a BioTek Synergy H1 plate reader. Crystal violet absorbance in all figures is expressed relative to absorbance in the *E. faecalis* monoculture biofilm wells.

### Determination of bacterial viability of biofilms

Biofilms were established in triplicate in tissue culture treated 24-well plates as described above and incubated for 24 hours at 37°C. Supernatants were removed, biofilms were gently washed with 1mL of sterile 1x PBS, and scraped as described above to resuspend. Suspensions were then serially diluted and plated onto appropriate agar using an EddyJet 2 spiral plater (Neutec Group) for determination of CFUs using a ProtoCOL 3 automated colony counter (Synbiosis).

### Growth curves

Overnight cultures of bacteria were adjusted to approximately 2x10^7^ CFU/mL in the various growth media described above. 200 µL of the adjusted bacterial suspension was distributed in at least triplicate wells of a clear 96-well plate and incubated in a BioTek Synergy H1 96-well plate reader at 37°C with continuous double-orbital shaking and a 1°C temperature differential between the top and bottom of the plate to prevent condensation. Bacterial growth was assessed via absorbance (OD_600_) every 15 minutes for a period of 18 hours. For assessment of CFUs, 5 mL bacterial suspensions were incubated at 37°C with shaking at 225 RPM, aliquots were taken hourly, serially diluted, and plated onto appropriate agar using an EddyJet 2 spiral plater (Neutec Group) for determination of CFUs using a ProtoCOL 3 automated colony counter (Synbiosis).

### Biofilm compositional analysis

Biofilms were established as described above in 24-well plates and incubated for 20 hours at 37°C, with the exception that an entire 24-well plate was used for each inoculum. Compositional analysis was performed as previously described for *P. mirabilis*^77^. Briefly, supernatants were removed and all wells of the entire 24-well plate were gently resuspended into a total volume of 3 mL of sterile Milli-Q water, an aliquot was removed from this total volume to generate the biofilm suspension fraction (BS). Suspensions were fixed for 1 hour with formaldehyde (37%) by incubating at room temperature with shaking at 200 RPM. 1 M NaOH was then added and samples were incubated for 3 hours at room temperature with shaking at 200 RPM. Samples were then centrifuged (20,000xg) for 1 hour at 4°C. Supernatant containing the soluble extrapolymeric substance (EPS) was removed and placed in a sterile microcentrifuge tube, while the remaining pellet was resuspended in 1 mL Milli-Q water to generate the cell fraction (CF). The EPS was filtered through a 0.22 µm filter and then was transferred to Slide-A-Lyzer dialysis cassette (Thermofisher, Cat# 66380) and placed in beaker containing Milli-Q water. The Milli-Q water was replaced twice after 2 hours, after which the sample was left to dialyze overnight. Samples were removed from the dialysis cassette to generate the EPS fraction (EPS). All samples were stored at -20C until end point analysis. Total eDNA was determined using the PicoGreen assay (Invitrogen, MP07581), total carbohydrate was determined using the Total Carbohydrate Assay Kit (Sigma, Cat# MAK104-1KT), and total protein was determined via Pierce BCA protein assay kit (Thermofisher, Cat# 23250), all following the manufacturer’s instructions ^77,78^. For comparison of wild-type and mutant polymicrobial biofilm protein content, biofilms were established in 24-well plates and grown for 24 hours at 37°C, after which three wells were scrapped, pooled, and analyzed as described above.

### Liquid-chromatography mass spectrometry (LC-MS) analysis of bacterial biofilms

Sample preparation and data analysis are described in detail in Supplemental Item 5. The mass spectrometry proteomics data have been deposited to the ProteomeXchange Consortium via the PRIDE partner repository with the dataset identifier PXD041693 ^79^.

### Mouse model of CAUTI

CAUTI studies were performed as previously described ^42,47,80^. In short, the inoculum was prepared by washing overnight cultures in PBS and adjusting to an OD_600_ of 0.2 for *P. mirabilis* and OD_600_ of 0.4 for *E. faecalis* (∼2x10^8^ CFU/mL), then diluting 1:100 to make a final inoculum of 2x10^6^ CFU/mL. Co-challenge inocula were generated by combining a 50:50 mix of each single-species inoculum. Female CBA/J mice aged 6-8 weeks (Jackson Laboratory) were anesthetized with a weight appropriate dose of ketamine/xylazine (80-120mg/kg ketamine and 5-10 mg/kg xylazine) via IP injection, after which mice were inoculated transurethrally with 50 µL of the appropriate inoculum suspension, delivering ∼1x10^5^ CFU/mouse. A 4 mm segment of sterile silicone tubing (0.64 mm O.D., 0.30 mm I.D., Braintree Scientific Inc.) was advanced into the bladder during inoculation and retained there for the duration of the study as done previously ^47,81^. After 96 hours, urine was collected, bladders, kidneys, and spleens were harvested and placed into 5 mL Eppendorf tubes containing 1 mL 1x PBS and 500 µL of 3.2mm stainless steel beads. Tissues were homogenized using a Bullet Blender 5 Gold (Next Advance, Speed 8, 4 minutes). Bladders were treated to two cycles to ensure full homogenization. Tissue homogenates were serially diluted and plated onto appropriate agar using an EddyJet 2 spiral plater (Neutec Group) for determination of CFUs using a ProtoCOL 3 automated colony counter (Synbiosis).

### Statistical analysis

Statistical significance of experimental results for biofilm, CFU, and growth curve data was assessed by two-way analysis of variance (ANOVA) multiple comparisons or one-way ANOVA as indicated in figure legends. For CAUTI model results, CFUs data was assessed by one-way ANOVA of log_10_ transformed data, and chi-square tests were used to analyzed incidences of abnormalities and health events. These analyses were performed using GraphPad Prism, version 9.3 (GraphPad Software) with a 95% confidence interval.

## Acknowledgments

We would like to thank Dr. Thomas Russo and members of his laboratory for helpful comments and critiques. This work was funded by the National Institute of Diabetes and Digestive and Kidney Diseases under award R01 DK123158 to CEA. The content is solely the responsibility of the authors and does not necessarily represent the official views of the National Institutes of Health.

## Author contributions

B.C.H. and C.E.A. designed experiments, analyzed data, and prepared the manuscript. B.C.H., V.B., J.V., L.B.G., S.M.T., B.S.L., and A.L.B. performed experiments. S.S. and J.Q. performed and analyzed the proteomics data. All authors reviewed the manuscript.

