## Supplemental Figures and Text for "Metabolic interplay between *Proteus mirabilis* and *Enterococcus faecalis* facilitates polymicrobial biofilm formation and invasive disease"

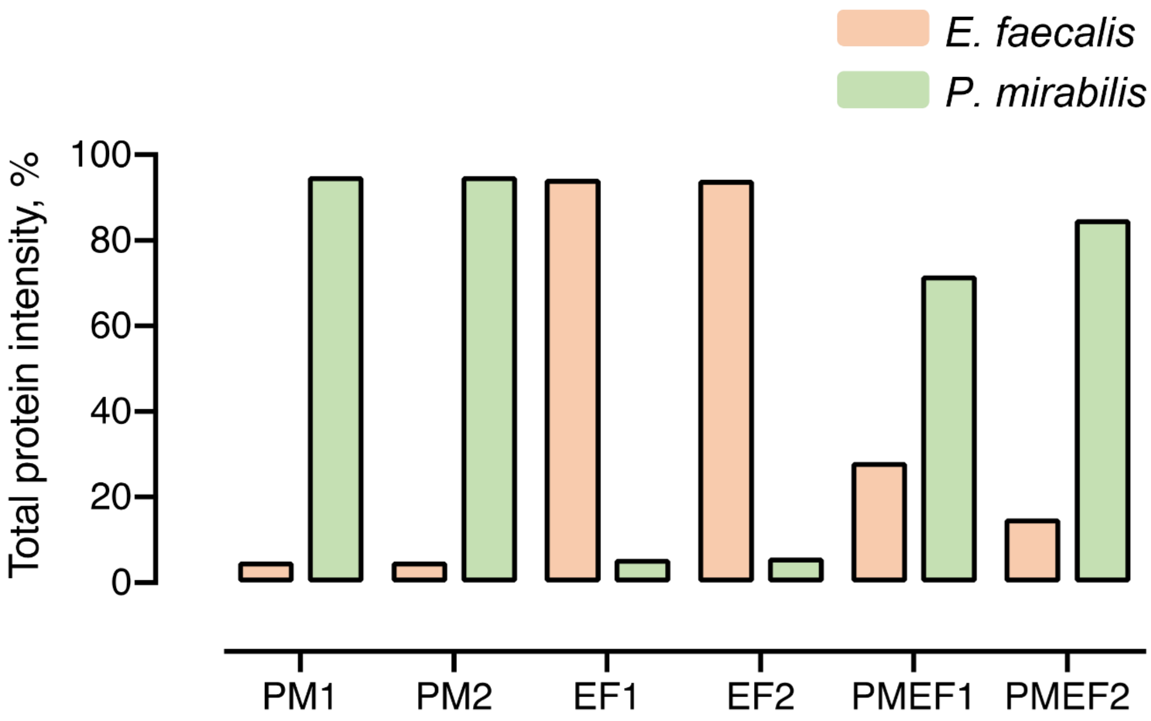


**Supplemental Figure 1. Total protein intensities for single and *P. mirabilis* and *E. faecalis* polymicrobial biofilms.** Protein identification was performed by searching against a combined database of *P. mirabilis* and *E. faecalis* protein sequence.

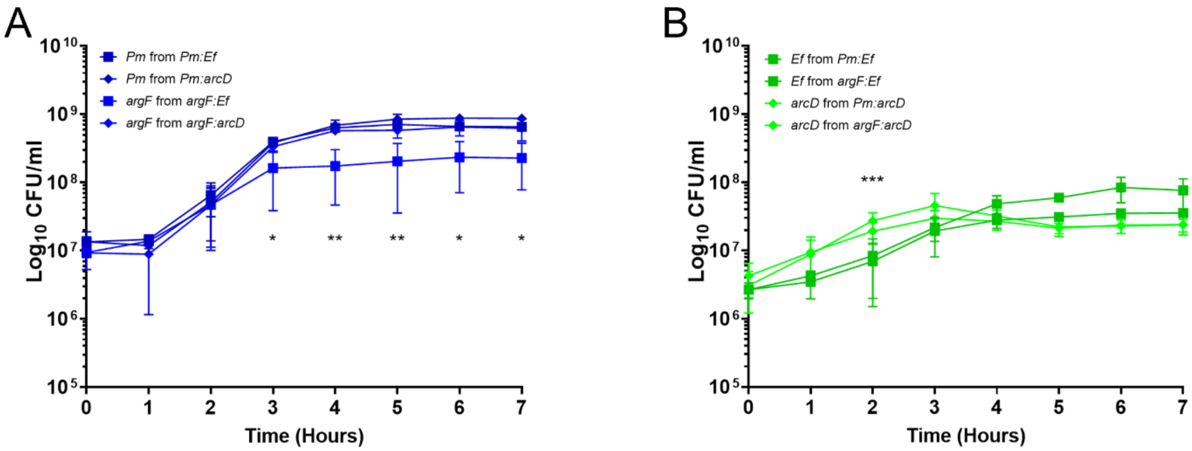


**Supplemental Figure 2. Fitness of *P. mirabilis argF, P. mirabilis speF,* and *E. faecalis arcD* during growth in human urine.** *P. mirabilis* and the *argF* mutant were co-cultured with either *E. faecalis* or the *arcD* mutant in human urine, and samples were plated every hour for determination of CFUs. A) *P. mirabilis* and B) *E. faecalis* CFU counts from the urine co-cultures. Error bars represent mean and standard deviation. *$P$<0.05, **$P$<0.01 by two-way ANOVA comparison of *argF* CFUs from *argF+Ef* to *argF* and wild-type CFUs from the other co-cultures in panel A, and for *arcD* CFUs compaed to *E. faecalis* CFUs in panel B.

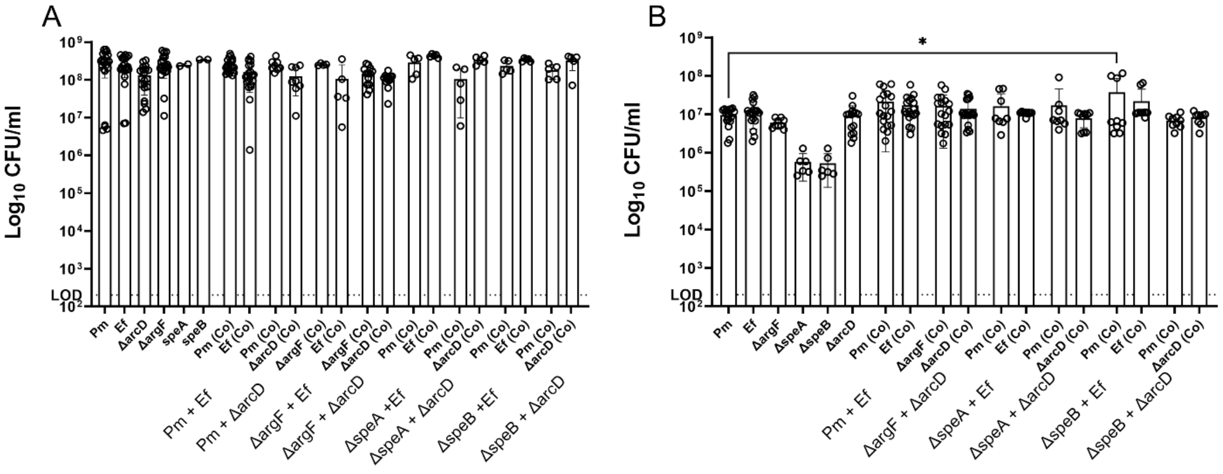


**Supplemental Figure 3. Changes in colony forming units between wild-type and mutant biofilms is insignificant and not a driving factor for the differences in biofilm biomass.** CFUs of biofilms grown for 24-hours in A) TSB-G or B) pooled human urine. Data represent the mean ± standard deviation for at least three independent experiments with at least two replicates each. ns = non-significant, * = P<.05 as determined by One-way ANOVA.

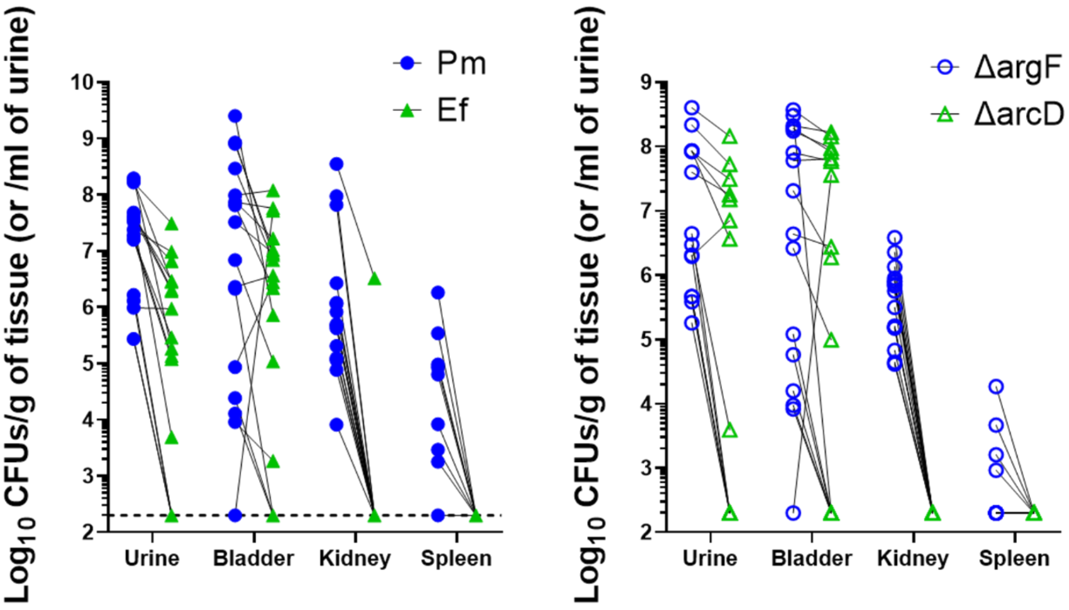


**Supplemental Figure 4. Colony forming units of each species from coinfected mice.** *P. mirabilis* and *E. faecalis* bacterial counts in urine (U), bladder (B), kidney (K), and spleen (S) homogenates. The CFUs from an individual coinfected mouse are connected with a black line for each organ.

**Supplemental Item 5. LCMS methodology and analysis.** *Protein digestion:* Biofilm suspension fraction (BS) was prepared as described above. After which, a surfactant-aided precipitation/on-pellet digestion method was adopted in the current study for sample preparation [82]. In brief, 100 µg protein was aliquoted from each sample and diluted to 1 µg/µL with 1% SDS. Protein was sequentially reduced by 10 mM dithiothreitol (DTT) at 56°C for 30 min and alkylated by 25 mM iodoacetamide (IAM) at 37°C in darkness for 30 min. Both steps were performed with rigorous vortexing in a thermomixer (Eppendorf). A total of 6 volumes of chilled acetone was then added to each sample with constant vortexing, and the mixture was incubated at -20°C for 3 hr. After centrifugation at 20,000 g, 4°C for 30 min, liquid was decanted, and protein pellet was gently washed by 500 µL methanol and air-dried for 1 min. A volume of 80 µL 50 mM pH 8.4 Tris-formic acid (FA) was then added, and samples were sonicated to loosen the protein pellet. A total volume of 20 µL trypsin (Sigma Aldrich, dissolved in 50 mM pH 8.4 Tris-FA) was added for 6-hr digestion at 37°C with rigorous vortexing in a thermomixer. Digestion was terminated by addition of 1 µL FA, and samples were centrifuged at 20,000 g, 4°C for 30 min. Supernatant was carefully transferred to LC vials for analysis.

*LC-MS analysis:* The LC-MS system consists of a Dionex µLtimate 3000 nano LC system, a Dinex µLtimate 3000 micro LC system with a WPS-3000 autosampler, and a ThermoFisher Orbitrap Fusion Lumos mass spectrometer. A large-inner diameter (i.d.) trapping column (300-um i.d. x 5 mm) was coupled to the nano LC column (75-um i.d. x 65 cm, packed with 2.5-um Xselect CSH C18 material) for high-capacity sample loading, cleanup and delivery. For each sample, 4 µL derived peptide was injected for LC-MS analysis. Mobile phase A and B were 0.1% FA in 2% acetonitrile (ACN) and 0.1% FA in 88% ACN. The 180-min LC gradient profile was: 4% for 3 min, 4–11 for 5 min, 11–32% B for 117 min, 32–50% B for 10 min, 50–97% B for 5 min, 97% B for 7 min, and then equilibrated to 4% for 27 min. The mass spectrometer was operated under data-dependent acquisition (DDA) mode with a maximal duty cycle of 3 s. MS1 spectra was acquired by Orbitrap (OT) under 120k resolution for ions within the m/z range of 400-1,500. Automatic Gain Control (AGC) and maximal injection time was set at 120% and 50 ms, and dynamic exclusion was set at 45 s, ± 10 ppm. Precursor ions were isolated by quadrupole using a m/z window of 1.2 Th, and were fragmented by high-energy collision dissociation (HCD). MS2 spectra was acquired OT under 15k resolution with a maximal injection time of 50 ms. Detailed LC-MS settings and relevant information are enclosed in a previous publication by Shen et al.[83].

*Data processing:* LC-MS files were searched against a NCBI protein sequence database containing both *Proteus mirabilis* and *Enterococcus faecalis* protein sequences using Sequest HT embedded in Proteome Discoverer 1.4 (ThermoFisher Scientific). Target-decoy searching approach using a concatenated forward and reverse protein sequence database was employed for global FDR estimation and control. Searching parameters include: 1) Precursor ion mass tolerance: 20 ppm; 2) Product ion mass tolerance: 0.02 Da; 3) Maximal missed cleavages per peptide: 2; 4) Fixed modifications: carbamidomethylation of cysteine; 5) Dynamic modifications: Oxidation of methionine, Acetylation of peptide N-terminals. Peptide filtering, protein inference and grouping, and FDR control were accomplished by Scaffold v5.0.0 (Proteome Software, Inc.) The filtered peptide-spectrum match (PSM) list was exported. Protein quantification was performed using IonStar, an in-house developed MS1 ion current-based quantitative proteomics method [84]. Peptide quantitative features were first generated by a two-step procedure encompassing 1) Chromatographic alignment with ChromAlign for inter-run calibration of retention time (RT) shift; ii) Data-independent MS1 feature generation a direct ion-current extraction (DICE) method, which extracts ion chromatograms for all precursor ions with corresponding MS2 scans in the aligned dataset with a defined m/z-RT window (10 ppm, 1 min). Both steps were accomplished in SIEVE v2.2 (ThermoFisher Scientific). Post-feature generation data processing was accomplished by UHR-IonStar v1.4 (https://github.com/JunQu-Lab/UHRIonStarApp) [85]. The filtered PSM list and the quantitative features database were first integrated by MS2 scan number to generate a list of annotated frames with peptide sequence assignment. The annotated frames were then subjected to dataset-wide normalization, principal component-based detection and removal of peptide outliers, and data aggregation to protein level. Protein quantification results were exported and manually curated and processed in Microsoft Excel.
